# Protein Function Prediction via Contig-Aware Multi-Level Feature Integration

**DOI:** 10.1101/2025.08.07.669053

**Authors:** Liang Yang, Kaixin Du, Yintong Lu, Mengzhu Wang, Hongxing Zhang, Shuo Yang, Yu Lin, Jiaming Zhuo, Duoyue Zhang, Yiqi Jiang, Xianglilan Zhang, Shuaicheng Li

## Abstract

Proteins play a central role in biological processes, and accurately predicting their functions is crucial for biomedical research. While computational methods have advanced significantly, most approaches rely solely on sequence or structure, neglecting critical inter-protein relationships, such as the topological arrangement of coding sequences (CDSs) within contigs. To address this gap, we propose CAML, a novel deep learning model that integrates intra-protein features including sequence and predicted structure with inter-protein features capturing functional linkages among CDSs in contigs. Specifically, CAML employs a Graph Isomorphism Network (GIN) to extract structural features from predicted protein contact graphs and ESM-2 for sequence embeddings. Additionally, it leverages kmer frequencies and a Bidirectional Long Short-Term Memory (BiLSTM) network to model functional relationships among colocalized CDSs within contigs, capturing operon-like associations. Extensive experiments demonstrate that CAML outperforms the state-of-the-art methods in accuracy, precision, recall and F1-score, achieving improvements of 11.24%, 12.43%, 13.59%, and 13.30%, respectively over the second-best model. Ablation studies further confirm the critical contribution of CAML’s multi-level biological feature integration in enhancing functional annotation accuracy. Our study demonstrates the importance of integrating structural, sequential, and CDSs topological features for accurate protein function prediction, providing a robust computational framework for genomics research.

## I. Introduction

Proteins execute essential biological processes, including immune response, signal transduction, and cellular communication[1]. Understanding their functions facilitates the elucidation of disease mechanisms, the development of therapeutics, and the advancement of precision medicine. The Clusters of Orthologous Groups (COG)[2] database classifies protein functions based on evolutionary conservation, organizing them into 26 functional categories such as Metabolism, Cellular Processes, and Information Storage. While the COG database provides a standardized classification system, its coverage remains limited. For instance, UniProtKB[3] contains over 200 million protein sequences, yet less than 0.1% of these have COG annotations. This limitation primarily results from its reliance on experimentally verified prokaryotic data. High-throughput sequencing further expands this disparity, as the rate of newly discovered protein sequences outpaces functional annotation efforts. This discrepancy highlights the need for computational approaches to efficiently and accurately predict protein functions using available biological data.

Computational methods for protein function prediction utilize diverse biological features, including protein sequences[4], domain architectures[5], family annotations[6], and protein-protein interaction (PPI) networks[7]. However, many proteins lack comprehensive feature annotations, restricting the applicability of these approaches. Due to this limitation, various sequence-only methods have been developed. For instance, BlastKNN[8] and Diamond[9] predict functions based on sequence similarity, but their accuracy declines when handling novel sequences with no detectable homology to annotated proteins. Recent advances in deep learning have facilitated the emergence of numerous deep learning-based methods for protein function annotation.

Several deep learning methods leverage Gene Ontology (GO) annotations to enhance protein function prediction. For instance, DeepGOPlus[10] integrates convolutional neural networks (CNNs) with sequence homology scores, while TALE[11] employs Transformer architectures to model sequence features alongside GO label dependencies. Other approaches, such as DeepGraphGO[12] and NetGO[13], further incorporate protein-protein interaction networks or multi-task learning to exploit GO hierarchies. It is important to note that these methods (DeepGOPlus, TALE, DeepGraphGO, and NetGO) are explicitly optimized for GO-based prediction; their high accuracy relies on GO-structured feature engineering and may not generalize to alternative annotation frameworks without significant modification.

Furthermore, other deep learning methods have begun to exploit the structure-function relationship, which is a well-established paradigm in molecular biology. Structural similarity often implies functional similarity, even in the absence of sequence conservation[14]. This principle underscores the importance of incorporating structural data to improve function prediction accuracy, particularly for sequences with low homology. DeepFRI[15] combines experimentally resolved structures with sequence embeddings to construct residue interaction graphs, which are then processed using graph convolutional networks (GCNs). The development of accurate structure prediction tools has further enabled structure-based approaches, as demonstrated by GAT-GO’s[16] use of graph attention networks on predicted structures. Nevertheless, existing structure-based methods remain constrained by their singular focus on individual protein features, overlooking potentially informative inter-protein relationships including conserved synteny arrangements of CDSs within contigs. Therefore, fully leveraging topological relationships of CDSs on contigs represents a critical direction for further advancements.

A contig represents a continuous DNA sequence derived from genome assembly[17]. Within the same contig, adjacent CDSs can exhibit functional linkages. This association arises from various biological mechanisms, including operon structures[18], metabolic gene clusters[19], and horizontal transfer units[20]. For instance, in prokaryotes, functionally related genes often co-localize as operons. Enzymes involved in specific metabolic pathways frequently cluster together, and resistance genes or mobile elements commonly exist as complete functional modules within a single contig. Therefore, the strategic integration of CDS positional topology within contigs for enhanced protein functional annotation represents an underexplored direction in computational genomics.

In this study, we introduce CAML, a protein function annotation model that integrates both intra-protein and inter-protein features (Fig. 1). Firstly, CAML takes protein sequences and their predicted structures as input. It utilizes a layer-wise GIN to extract structural features and employs ESM-2 to derive sequence features, which serve as node features for the contact graph. Then, to leverage the functional similarity of CDSs within the same contig and further improve prediction, the model employs a k-mer frequency-based approach for extracting nucleotide sequence features from contigs. Simultaneously, a BiLSTM captures the functional topological relationships among co-localized CDSs. Experimental results demonstrate that the incorporation of protein structure, sequence, and CDS topological relationships from contigs leads to enhanced protein function prediction accuracy. Compared to state-of-the-art methods, CAML achieves significant improvements across multiple metrics: accuracy (11.24% ~56.64%), precision (12.43%~ 46.18%), recall (13.59% ~62.72%), and F1-score (13.30% ~60.84%). Notably, CAML is database-independent, unlike GO-specialized deep learning methods that rely on prior knowledge of specific databases; our model can seamlessly generalize from COG to GO annotations by inherently learning functional principles rather than relying on predefined database ontologies. To our knowledge, CAML is the first tool to integrate contig topology into protein function prediction, unlocking novel biological insights for co-localized CDSs. By combining multimodal protein features with genomic context, CAML enables precise protein function annotation, offering a generalizable and high-performance solution for genomics research.

**Fig. 1.**
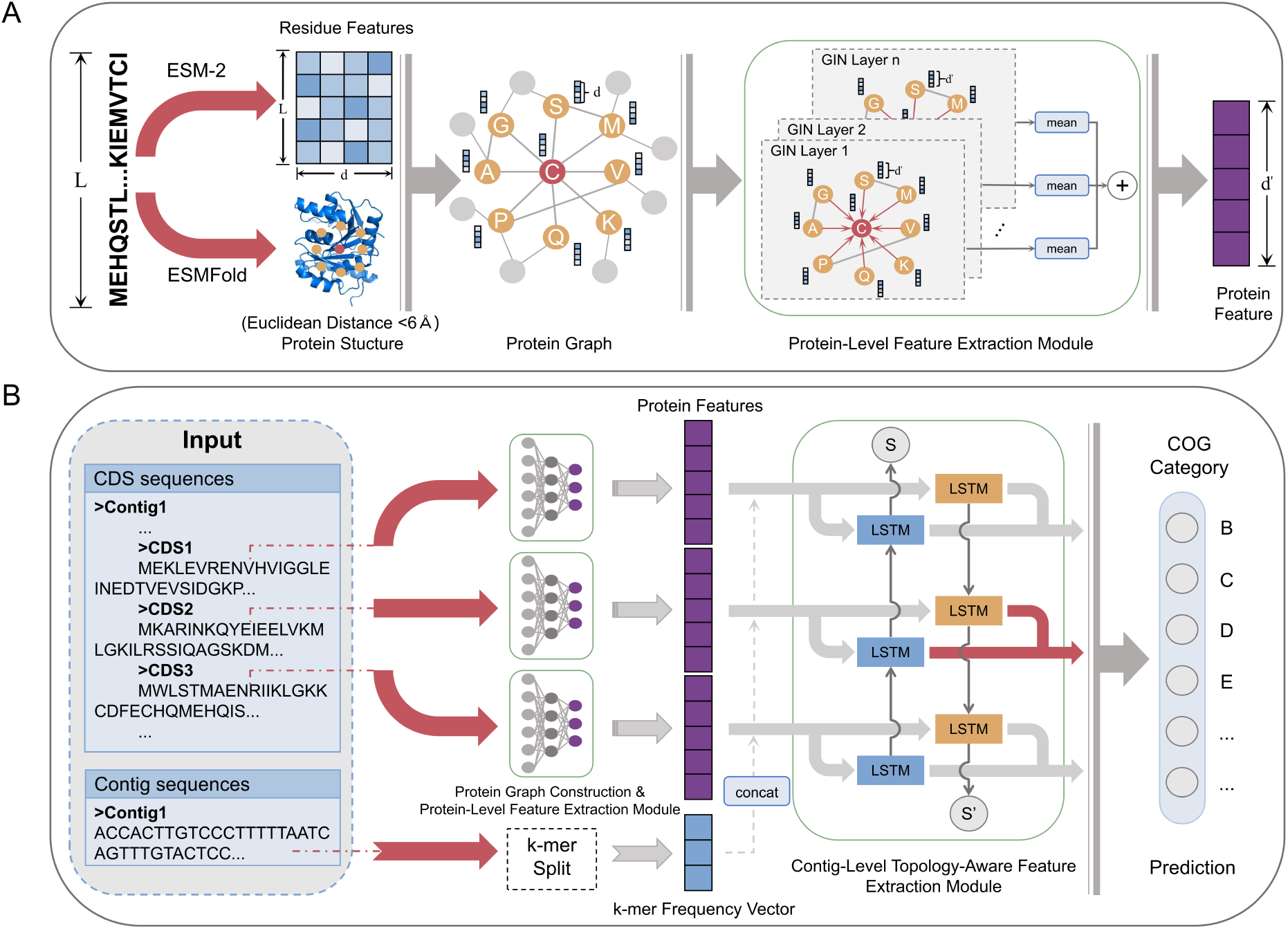
Overall architecture of CAML. **(A)** The protein sequence is initially processed by ESMFold and ESM-2 to generate its structure and residue-level features. Based on the protein structure and residue features, a protein graph is constructed. This graph is then fed into multiple layers of GIN to propagate residue features. Finally, the residue features from each GIN layer are aggregated via mean pooling and concatenated to form the protein’s final feature representation. **(B)** To achieve more accurate protein function annotation, all CDS sequences on a contig are used as input. These sequences are processed through protein graph construction and protein-level feature extraction module to obtain their corresponding protein features, which are then concatenated with the k-mer frequency vector of the contig’s nucleotide sequence. After arranging the protein features in the same order as the CDS positions on the contig, they are divided into protein feature fragments of length 3 using a sliding window approach, which are then fed into a BiLSTM to capture topological information. The BiLSTM’s output is then used for the final prediction of protein functions.

## II. Methodology

### A. Overview

As shown in Fig. 1, The model construction process consists of four main modules: (i) Protein graph construction. This module takes protein sequences as input, extracts sequence features using ESM-2, and predicts protein structures using ESMFold. It constructs a contact graph from the protein structure and uses the sequence features as residue-level features. (ii) Protein-level feature extraction. This module propagates the node features of the protein graph through N layers of GIN layers, and finally combines the graph-level features from each GIN layer as the protein features. (iii) Protein feature enhancement. This module takes the protein features corresponding to the CDS on the same contig and the contig sequence as input. The contig sequence is split into k-mers to obtain a k-mer frequency vector, which is then concatenated with the protein features to generate enhanced protein features. (iv) Contig-level topology-aware feature extraction. This module takes the enhanced protein features as input and employs a BiLSTM to perform protein feature aggregation, thereby obtaining topology-aware protein features by modeling the relationships among proteins encoded on the same contig.

### B. Protein graph construction

ESM-2 is an upgraded protein language model from Meta AI[21], using deeper transformers and improved self-supervised learning on larger datasets than ESM-1b[22]. ESM-2 demonstrates enhanced capabilities in capturing fundamental biological principles of proteins, achieving state-of-the-art performance across diverse downstream applications. With pre-trained ESM-2, we can extract comprehensive and accurate information embedded in protein sequences, including evolutionary relationships and structural insights. ESMFold is a fast structure prediction tool based on the protein language model ESM-2. It predicts 3D structures directly from single sequences using a pre-trained attention mechanism without requiring multiple sequence alignments. ESMFold achieves both accuracy and efficiency, making it suitable for high-throughput prediction.

We employ the esm2 t30 150M UR50D model from ESM-2 to derive protein sequence features. This model processes input sequences through 30 transformer layers, with each successive layer building upon the previous one to produce increasingly complex embeddings that capture higher-level evolutionary patterns. The 30th-layer(final) output best captures comprehensive sequence information through hierarchical refinement, thus serving as our final protein sequence features *x*^*esm*2^. For each protein, we use the esmfold_v1 model from ESMFold to predict protein structures *x*^*esmfold*^. Subsequently, we build the protein contact graph from structure *x*^*esmfold*^, where sequence features *x*^*esm*2^ serves as the residue-level node features. Let *G*(*V, E*) represent the contact graph of protein, where *v*_*i*_ ∈ *V* denotes a residue *r*_*i*_ from structure *x*^*esmfold*^ and *e*_*i,j*_ ∈ *E* represents an edge connecting residue pairs (*v*_*i*_, *v*_*j*_). An edge *e*_*i,j*_ exists if and only if the Euclidean distance between the *C*_*α*_ atoms of two residues *r*_*i*_, *r*_*j*_ in structure *x*^*esmfold*^ is less than 6 Å.

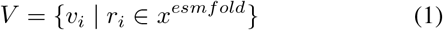

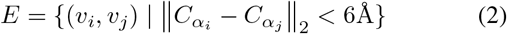

Next, for each residue *r*_*i*_, its sequence feature 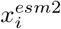 is assigned as the initial feature of the corresponding node *v*_*i*_.

### C. Protein-level feature extraction

In this module, the model takes the constructed protein graph *G*_*p*_(*V*_*p*_, *E*_*p*_) of protein *p* as input and propagates node features through N layers of GIN to obtain more structure-aware node features.

For the (*l* + 1)-th GIN layer, the feature update process of node *v* ∈ *V*_*p*_ is as follows:

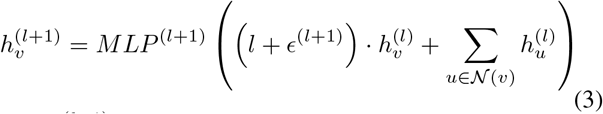

where 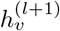 represents the output feature of node *v* at the (*l* + 1)-th layer, 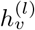 is the feature of node *v* at the *l*-th layer, *MLP* ^(*l*+1)^ is the multilayer perceptron(MLP) of the (*l* + 1)-th layer, *ϵ*^(*l*+1)^ is a learnable parameter of the (*l* + 1)-th layer, and 𝒩(*v*) is the set of 1-hop neighbor nodes of node *v*.

The node features from each GIN layer are aggregated via mean pooling to generate layer-wise protein features *h*_*p*_ of protein *p*. To capture diverse structural views, we concatenate all layer-wise protein features and pass them through an MLP to obtain the final protein features 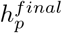.

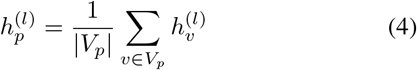

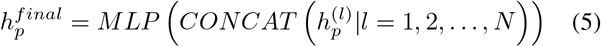

Where 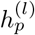 denotes the protein-level feature generated by mean pooling at layer *l, V*_*p*_ is the collection of all nodes in the protein graph *G*_p_, |*V*_p_ | is the number of nodes, and 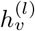 is the feature of node *v* at the *l*-th layer.

### D. Protein feature enhancement

A k-mer is a contiguous substring of length k in nucleotide sequences or amino acid sequences, serving as a fundamental unit in bioinformatics analysis[23]. By splitting sequences into overlapping k-mers, it quantifies local sequence features. We encode the contig’s nucleotide sequence into a k-mer frequency vector to characterize its sequence features.

Given a nucleotide sequence *S* = *s*_1_*s*_2_…*s*_*L*_ of length *L*, where each *s*_*i*_ ∈ {*A, T, C, G*}, we extract all possible k-mers via a sliding window approach. Specifically, the k-mer *w*_*i*_ starting at position *i* is defined as:

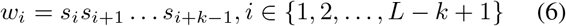

This process generates *N* = *L* − *k* + 1 overlapping k-mers. All possible k-mers are lexicographically ordered into a set 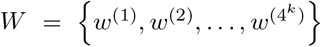, where the total number of unique k-mers is 4^*k*^. The k-mer frequency vector 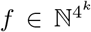 is constructed such that its *j*-th component *f*_*j*_ counts the occurrences of the k-mer *w*^(*j*)^ in *S*.

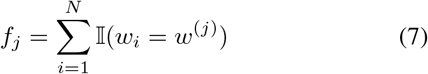

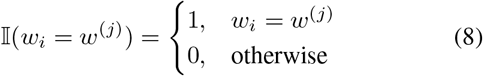

Then, the k-mer frequency vector is represented as 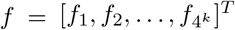. To eliminate the bias of sequence length, the frequency vector is divided by the total number of k-mers *N*. It is then normalized to a range of [0,1] through a normalization layer. Subsequently, the normalized frequency vector is input into an MLP to obtain the final k-mer frequency vector *f* ^*k*−*mer*^.

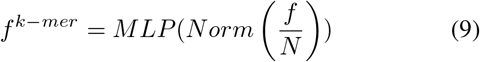

To further enhance the obtained protein feature 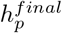 generated in the previous module and fully leverage the functional correlations among proteins located on the same contig *C*, we concatenate the protein feature 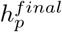 with a 3-mer frequency vector 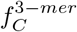 derived from the nucleotide sequence of the contig *C*.

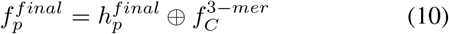

where 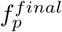 is the final enhanced protein feature of protein *p*, ⊕ denotes the concatenation operator.

### E. Contig-level topology-aware feature extraction

To capture the functional and topological relationships among proteins encoded by the same contig, we organize their feature vectors according to genomic coordinates to construct a protein feature sequence:

Given *n* CDSs on a contig with corresponding protein feature vectors 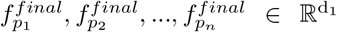, the input sequence matrix is constructed as:

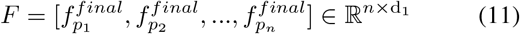

Then, it is divided into *m* segments using a sliding window of size 3, and ensure that the central protein feature’s corresponding CDS is labeled. The *i*-th segment can be represented as:

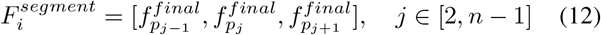

The segmented triplet matrices can be represented as 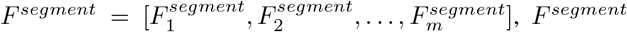 is then fed into a BiLSTM to aggregate protein features across each triplet.

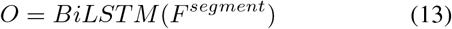

The output of the BiLSTM 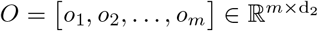 is the central CDS feature of each triplet. Then *O* is input into the final MLP prediction layer:

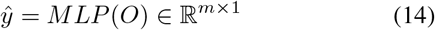

In this study, we use cross entropy as the loss function:

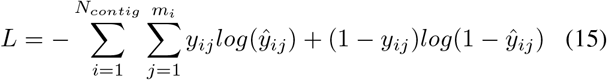

where *N*_*contig*_ is the total number of contigs, *m*_*i*_ is the total number of labeled CDSs on the *i*-th contig, *y*_*ij*_ is the ground truth of the central CDS of the *j*-th triplet on the *i*-th contig, *ŷ*_*ij*_ is the predicted probability value of the central CDS of the *j*-th triplet on the *i*-th contig.

## III. Experiments

### A. Datasets

We downloaded assembly data of the human body from the MGnify[24] database, which is an EMBL-EBI microbiome database offering standardized storage, functional annotation, and interactive analysis of metagenomic data. We downloaded assembly data comprising 15,000 contigs, including nucleotide sequences of the contigs and amino acid sequences of the CDSs. We subsequently applied FastANI[25] to remove contigs with ANI (Average Nucleotide Identity) similarity*>*85%, followed by CD-HIT[26] clustering to eliminate CDSs exhibiting*>*85% sequence identity. We additionally imposed a maximum length limit of 1,022 amino acids for CDS sequences to ensure all sequence features could be generated using ESM-2. To annotate CDSs with COG categories, we downloaded the COGorg24.faa.gz file from the NCBI COG database and performed sequence alignment using DIAMOND to assign COG classifications.

From the final processed dataset, we randomly selected 9,000 contigs for training, 1,000 contigs for validation, and 2,000 contigs for testing. The training set contained 58,803 CDSs including 28,136 labeled CDSs, while the validation and test sets comprised 6,779 CDSs with 3,139 labeled CDSs and 13,019 CDSs with 6,229 labeled CDSs respectively. Notably, only labeled CDSs were used to compute the model’s loss function, while unannotated CDSs were treated as part of the contig’s CDS sequence input to the BiLSTM. The details of benchmark dataset are described in Table I.

**TABLE 1.** Details on benchmark dataset.

| Genomic Component | Training Set | Validation Set | Test Set |
| --- | --- | --- | --- |
| Contig | 9000 | 1000 | 2000 |
| CDS | 58803 | 6779 | 13019 |
| Labeled CDS | 28136 | 3139 | 6229 |

Fig. 2 illustrates the distribution of proteins across COG categories. The final dataset comprises 23 COG categories (categories A and Y were excluded due to absence of training samples, while category S was removed because of its functional unknown), demonstrating an imbalanced distribution: categories G and E each contain over 3,000 proteins, while categories X and B each contain fewer than 200 proteins (these counts represent the combined total across the training, validation, and test sets, the same applies hereinafter). From the perspective of protein proportions within each category, both the training & validation set and the test set exhibit imbalanced distributions. Furthermore, the proportion of each category’s proteins relative to the entire training & validation set or test set remains largely consistent between these two datasets. To facilitate experimental analysis, we further classified the categories into high-abundance(*>*2900 proteins), medium-abundance(600 ~ 2900 proteins), and low-abundance(*<*600 proteins) groups based on their respective protein counts.

**Fig. 2.**
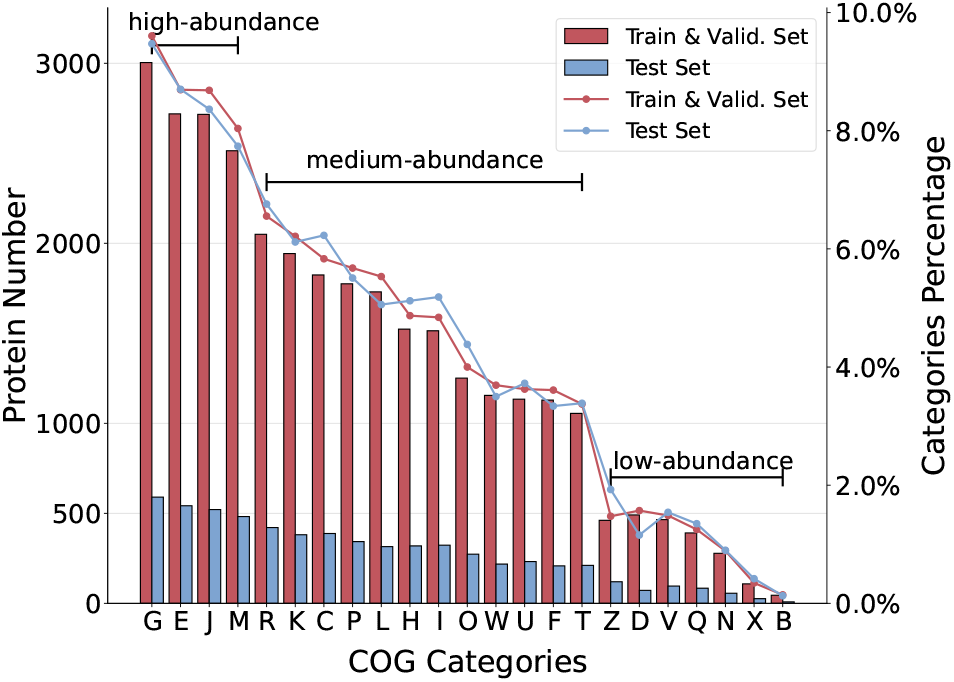
Category distribution of the dataset. The bar chart displays the number of proteins across different categories, while the two overlaid line graphs represent the relative proportions of proteins in the train & validation set and the test set, respectively. Based on protein abundance levels, the categories are further classified into high-abundance, medium-abundance, and low-abundance groups.

### B. Baselines

- DeepGOCNN[10]: takes protein sequences as input and employs pre-trained parameter-free one-hot encoding combined with multi-scale CNNs to extract motif features.
- DeepFam[27]: takes protein sequences as input, encodes the sequences using a custom one-hot encoding scheme, and then feeds them into multi-scale CNNs to extract motif features.
- DeepFRI: represents proteins as graphs and initializes node features using the pre-trained language model, called LSTM-LM.
- Struct2GO[28]: constructs a graph from AlphaFold-predicted protein structures and employs the Node2Vec graph representation learning method to initialize node features. Simultaneously, it integrates the SeqVec pre-trained model to capture semantic information from protein sequences.

### C. Evaluation Metrics

We evaluate these models using four evaluation metrics, including accuracy, precision, recall and F1-score. Accuracy measures the overall proportion of correct predictions. Precision quantifies the reliability of positive predictions. Recall assesses the model’s ability to identify all relevant instances. The F1-score, as the harmonic mean of precision and recall, balances these two metrics and is particularly useful in the case of unbalanced data.

### D. Overall Performance Evaluation

To evaluate the predictive performance of CAML, we compare it with four state-of-the-art methods: DeepGOCNN, DeepFam, DeepFRI, Struct2GO, and all these methods use the parameters with the best results.

Table II shows the experiment results. It can be seen that: (i) Obviously, the CAML achieves the best performance across all evaluation metrics. In terms of accuracy, precision, recall, and F1-score, the CAML achieved performance improvements of 11.24% ~56.64%, 12.43% ~46.18%, 13.59% ~62.72%, and 13.30% ~ 60.84%, respectively. (ii) Compared to structure-based models(DeepFRI and Struct2GO), the CAML demonstrates even greater superiority over sequence-based models(DeepGOCNN and DeepFam), which fully validates the effectiveness of incorporating protein structure into protein function annotation models. (iii) The CAML outperforms DeepFRI, which also utilizes protein structural information, primarily because DeepFRI does not leverage semantically richer pre-trained models(e.g., SeqVec, ESM-1b, or ESM-2) features to enhance protein function annotation. (iv) Compared to the second-best performing model, Struct2GO, the CAML achieved a significant lead, this is because Struct2GO did not utilize the semantically richer pre-trained SeqVec features for initializing the protein graph’s node features, but instead only employed them as sequence features, failing to fully capture the topological characteristics of protein structures. This is also because Struct2GO did not employ a GIN protein graph feature aggregation method to capture protein features from different structural views. More importantly, it failed to utilize the functional similarity characteristics of CDS on contigs to further enhance annotation accuracy.

**TABLE 2.**
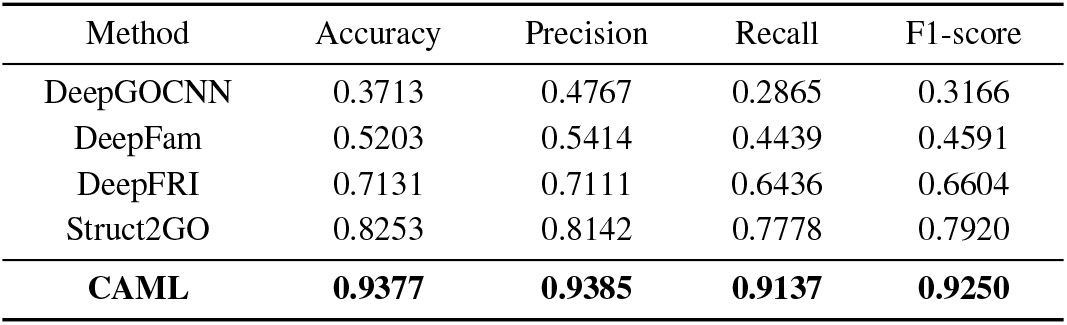
Comparative performance of different approaches with our proposed method. Best results are highlighted in **boldface**.

### E. Category-wise Performance Analysis

To validate CAML’s prediction performance across different categories and evaluate its robustness on datasets with imbalanced protein distributions, we conducted comprehensive category-level comparisons with baseline methods in terms of precision, recall, and F1-score (for Accuracy, the per-category calculation yields the same value as recall). Additionally, we employed a confusion matrix to perform detailed analysis of CAML’s classification accuracy.

As shown in Fig. 3, CAML demonstrated superior performance across nearly all categories, with only one exception: in category U, its precision was marginally lower (by 1.73%) than Struct2GO. The benchmarking results revealed a clear performance hierarchy: sequence-based models (DeepGOCNN and DeepFam) showed the lowest precision, recall and F1-score, while structure-aware methods (DeepFRI and Struct2GO) achieved progressively better results. CAML’s outstanding performance stems from its innovative integration of ESM-2 initialized graph features, GIN-based multi-view aggregation, and CDS functional similarity.

**Fig. 3.**
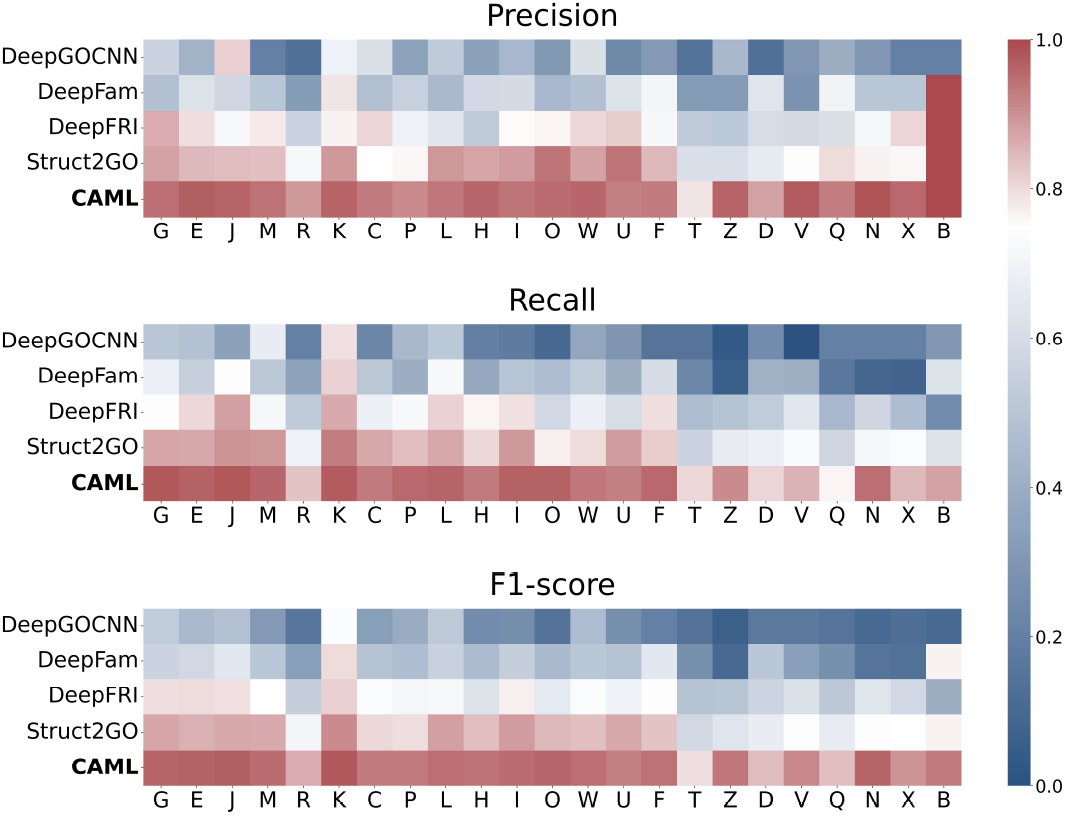
Comparative performance of different approaches with our proposed approach across 23 functional categories. Evaluation metrics are calculated separately for each category.

Performance analysis by abundance levels showed: (i) In high-abundance categories such as G and E, CAML achieved exceptional F1-scores of 0.9625 and 0.9676 respectively, representing improvements of 8.81%~ 43.26% and 11.15% ~51.88% over baselines. (ii) For medium-abundance categories, including O and W, CAML attained F1-scores of 0.9601 and 0.9466, outperforming baselines by 11.26% ~81.47% and 11.01% ~48.68%. (iii) Notably in low-abundance categories such as X and B, CAML maintained robust performance with F1-scores of 0.8980 and 0.9333, exceeding baselines by 15.29% ~77.50% and 16.41% ~83.33%. This comprehensive evaluation demonstrates CAML’s exceptional capability in protein function prediction, particularly its remarkable performance in few-shot learning scenarios with low-abundance categories.

Fig. 4 displays the confusion matrix of our CAML model across 23 COG categories. While most categories achieve high prediction accuracy, one subtle yet consistent patterns emerge: R-class (General function prediction only) proteins exhibit a slightly elevated misclassification rate, as reflected in its F1-score of 0.8601 compared to the overall average of 0.9250. This is particularly noticeable with functionally related categories like G-class (Carbohydrate transport and metabolism). This phenomenon likely stems from the functional ambiguity of R-class proteins. While their exact functions remain unknown, they often participate in fundamental cellular processes that overlap with or are regulated by pathways such as carbohydrate metabolism. These inherent similarities pose significant challenges for the classification model, leading to unavoidable performance degradation (e.g., Struct2GO’s R-class F1-score=0.7047 vs. overall 0.7920).

**Fig. 4.**
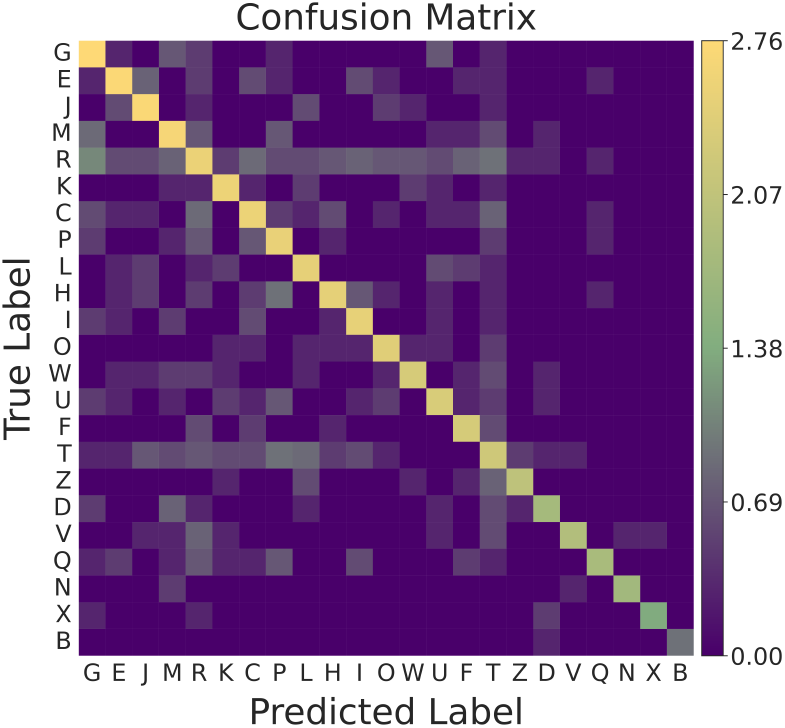
Confusion matrix of CAML across 23 functional categories on the test set, displaying log_10_(counts+1) values where diagonal entries represent correct predictions and off-diagonal entries indicate classification errors.

Collectively, these results demonstrate that CAML establishes state-of-the-art protein function prediction performance across all abundance categories. Its innovative combination of sequence-structure features and inter-protein CDS functional linkages proves particularly effective in achieving robust performance on imbalanced datasets.

### F. Ablation Analysis

To systematically evaluate the contributions of CAML’s key innovations to its performance improvement, we conducted a comprehensive ablation study with three components: (i) To validate the effectiveness of initializing protein graph node features with ESM-2 sequence features, we replaced the ESM-2 sequence features in CAML with the SeqVec sequence features used in Struct2GO (CAML-SeqVec) and with the LSTM-LM sequence features used in DeepFRI (CAML-LSTM-LM). (ii) To evaluate the capability of the GIN protein graph feature aggregation algorithm, we replaced it with GCN and GAT in CAML(CAML-GCN, CAML-GAT). (iii) To evaluate the contribution of contig topological information modules in protein function annotation, we removed the BiLSTM-based topology-aware feature extraction module (CAML-w/o BiLSTM), as well as all contig topological information, including the k-mer frequency vector and the BiLSTM-based topology-aware feature extraction module (CAML-w/o Contig).

The overall performance of these models is shown in Fig. 5(a). Based on the results, we can find that: (i) CAML demonstrates significant improvements over CAML-LSTM-LM across all evaluation metrics, highlighting the crucial role of state-of-the-art pretrained protein language model features in initializing protein graph nodes; furthermore, CAML’s superior performance over CAML-SeqVec indicates that ESM-2 features (640-dimensional) provide more comprehensive protein representation while being more lightweight than SeqVec features (1024-dimensional) for initializing protein graph node features. (ii) When compared with CAML-GAT/GCN, CAML shows consistent improvements in all metrics, demonstrating that the GIN-based graph feature aggregation algorithm effectively enhances function annotation through multi-structural view information. (iii) The progressive performance gains observed across CAML-w/o Contig, CAML-w/o BiLSTM, and CAML underscore the combined importance of k-mer frequency vectors and BiLSTM-based topology-aware feature extraction, confirming that both contig sequence features and CDS topological relationships are essential for accurate protein function annotation.

**Fig. 5.**
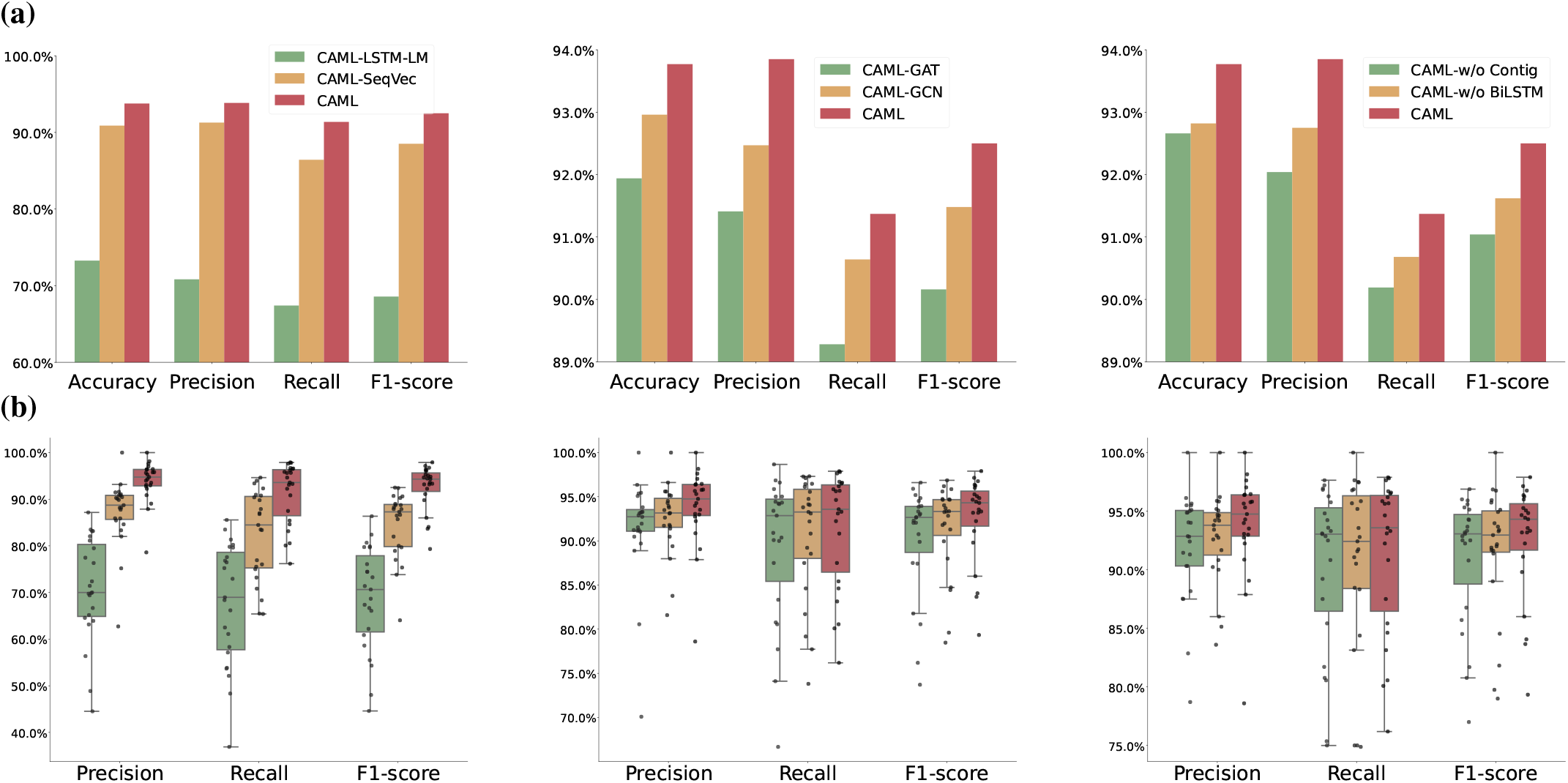
Comparison of performance between our method CAML and its variant for ablation evaluation. **(a)** Overall performance comparison. **(b)** Performance comparison across 23 functional categories. Evaluation metrics are calculated separately for each category. Each point represents a category. The median is represented by the centerline of the boxplot, while the first (Q1) and third (Q3) quartiles are indicated by the bounds of the box. The height of the box corresponds to the interquartile range (IQR=Q3−Q1). The whiskers represent the Q1−1.5*×*IQR and Q3+1.5*×*IQR, respectively. Since accuracy and recall yield identical values when computed per category, accuracy is not visualized.

Additionally, the performance of CAML and its variant models across different categories is illustrated in Figure 5(b). The results demonstrate that: (i) CAML, utilizing ESM-2 embeddings to initialize the node features of the protein graph, exhibits narrower interquartile ranges and superior median values across all metrics compared to CAML-LSTM-LM and CAML-SeqVec, indicating stable and outstanding prediction performance across all categories. Notably, CAML achieves median F1-score improvements of 23.67% and 7.02% over these variants, respectively. These results unequivocally confirm the unparalleled value of ESM-2-initialized protein structure graphs for protein function prediction. (ii) CAML employing the GIN-based graph feature aggregation algorithm demonstrates improved median values across all evaluation metrics compared to both CAML-GAT and CAML-GCN. Furthermore, it exhibits narrower interquartile ranges in F1-score distributions, providing evidence that the GIN-based algorithm effectively enhances protein function annotation through its integration of multi-structural view information. (iii) CAML with BiLSTM for capturing contig topological information demonstrates superior median values across all evaluation metrics compared to CAML-w/o BiLSTM. Although showing slightly wider interquartile ranges in recall, CAML maintains comparable interquartile ranges in F1-score (the harmonic mean of precision and recall), confirming the effectiveness of the BiLSTM-based topology-aware feature extraction module for accurate protein function annotation. CAML-w/o BiLSTM, which employs k-mer frequency vectors, achieves both superior median precision and higher overall F1-scores compared to CAML-w/o Contig, confirming the significant contribution of contig sequence features to accurate protein function annotation.

## IV. Discussion

In this study, we present CAML, a novel framework for protein function prediction that integrates intra-protein (sequence and structure) and inter-protein (CDS topological relationships on contigs) features. By initializing protein graph nodes with ESM-2 embeddings and employing layer-wise GIN aggregation, CAML achieves a comprehensive representation of protein features, unifying sequence and structural information. Furthermore, the incorporation of contig-level CDS topology via BiLSTM captures functional correlations among co-localized genes, a previously underexplored dimension in protein annotation.

Our experiments demonstrate CAML’s superiority over state-of-the-art methods, with significant gains in accuracy (up to 56.64%), F1-score (up to 60.84%), and other metrics. The category-wise performance analysis reveals that CAML not only delivers superior prediction accuracy across all abundance categories but also exhibits remarkable robustness when handling imbalanced datasets. Ablation studies validate the contributions of (i) leveraging ESM-2 embeddings for protein graph node initialization, which enables structure-aware representation learning unlike conventional sequence-only applications, (ii) GIN-based structural aggregation, and (iii) contig-derived CDS topology, representing a novel contribution in protein function prediction.

Beyond performance, CAML’s database-agnostic design represents a paradigm shift. Unlike GO-specific tools constrained by prior knowledge, CAML can seamlessly generalizes across annotation systems (e.g., from COG to GO), bridging the gap between genomic context and protein function. By unifying multimodal protein features with contig-level functional topology, CAML not only advances prediction accuracy but also unlocks new biological insights into co-localized CDSs. Future work will extend this framework to integrate multi-view features from diverse pre-trained models, further advancing the interpretability and scope of protein annotation. Ultimately, CAML learns the intricate relationships between protein structure, sequence, and genomic architecture, offering a universal and context-aware solution for next-generation functional genomics.

## ACKNOWLEDGMENTS

This work was supported in part by the National Natural Science Foundation of China (No. 62376088, 62406100), in part by the Hebei Natural Science Foundation (No. F2024202047), in part by the Hebei Yanzhao Golden Platform Talent Gathering Programme Core Talent Project (Education Platform)(HJZD202509), in part by the Tianjin Natural Science Foundation(No. 24JCQNJC00320), in part by the Young Collaborative Research Grant (C2004-23Y), and in part by the Beijing Postdoctoral Research Foundation.

